# Ecdysone Antagonizes Growth and Juvenile Hormone-Dependent Soldier Caste Identity in *Camponotus floridanus*

**DOI:** 10.64898/2026.10.26.761948

**Authors:** Olivia MacMillan, Alexander Craig, Camryn Giddens, Julia Singer, David Qiu, Rajendhran Rajakumar

## Abstract

From tardigrades to lobsters, ecdysone is the critical steroid hormone regulating ecdysis across Panarthropoda. In insects, the sesquiterpenoid hormone called juvenile hormone (JH) prolongs the juvenile state, and together, ecdysone and JH regulate growth and developmental timing. The complex worker caste system in ants has elaborated multiple times and is composed of individuals exhibiting an array of size and allometry variation. While we have accumulated a great deal of knowledge related to the role of JH in influencing size and caste identity in ants, the role of ecdysone in caste development remains poorly understood. Here we show in *Camponotus floridanus* that ecdysone negatively regulates size, can block soldier determination, and there is an antagonistic relationship between JH and ecdysone. First, we show that during the worker-soldier switchpoint sensitivity window of development, ecdysone is capable of negatively influencing head size, body size, head-to-body allometry, and unexpectedly extends developmental timing. We then show at the molecular level that as expected, developing soldiers upregulate ecdysone degradation, yet surprisingly, developing soldiers also upregulate ecdysone synthesis and the reception of ecdysone signalling. We next investigated the interaction between JH and ecdysone signaling and found that the two hormones antagonize each other at both the molecular and phenotypic levels. Furthermore, in contrast to other insects, we uncovered that ecdysone signaling is negatively regulated by temperature. Beyond its role in size regulation, we also provide preliminary data suggesting that the interaction of JH and ecdysone may have sex inducing properties, generating male-like anomalies. Altogether, our findings show that ecdysone blocks JH-dependent soldier development and we propose that the dynamic regulation of ecdysone synthesis, reception, and degradation may be the basis for the complex sigmoid-like head-to-body allometry found in this species as well as several others across ants. More generally, the evolutionary tinkering of antagonistic growth hormones, their regulation and interaction, and opposing responses to environmental cues, may be a major source for size diversity across developmentally plastic organisms.

## Introduction

Since the Ediacaran over half a billion years ago, ecdysone has been the master regulator of ecdysis across the Panarthropoda (Rota-Stabelli et al., 2013; Schumann et al., 2018). Beyond ecdysis, ecdysone has been implicated in the (re-)formation of an array of traits including eyespots in butterflies, claw regeneration in fiddler crabs, and wing polyphenism in aphids (Brakefield et al., 1998; Das & Durica, 2013; Vellichirammal et al., 2017). In holometabolous insects, ecdysone is essential for the induction of metamorphosis and the sesquiterpenoid hormone, juvenile hormone (JH) prolongs development (Jindra et al., 2013; K. Li et al., 2019; Nijhout, 2022; Yamanaka et al., 2013). Furthermore, these two hormones antagonize each other (S. Liu et al., 2018). For example, in the fly *Drosophila melanogaster* and the silkworm *Bombyx mori*, JH increases the expression of the gene that encodes Krüppel homolog 1 (*Kr-h1*), the vital downstream transcription factor of the JH pathway, in the ecdysone-producing prothoracic gland (PG) (Zhang et al., 2018). This subsequently leads to a downregulation of the expression and function of the ecdysone synthesis enzymes in the PG (Zhang et al., 2018).

Ants live in a societies and individuals among their colony exhibit an array of phenotypic variation. This variation is optimized for the division of labour and found across a complex system of castes where large, winged queens conduct reproductive tasks and small workers conduct non-reproductive tasks. (Hölldobler & Wilson, 1990; Wheeler, 1991, 1991; Wheeler & Nijhout, 1981). Several genera of ants subdivide their worker castes into minors and soldiers, where minors are smaller and participate in brood care and foraging, whereas, soldiers are larger with disproportionately larger heads, and are imperative for the colony defense (Rajakumar et al., 2018; Wheeler, 1991; Wilson, 1953). These dramatically different morphologies are the product of developmental plasticity, where the environment acts on developmental processes giving rise to the production of the different castes (Simpson et al., 2011). Environmental variation involved in caste development include temperature (MacMillan et al., 2025; Passera, 1974) and nutrition, (Goetsch, 1937; Passera, 1974; Smith, 1942, 1944), and are modulated by epigenetic and hormonal mechanisms (Alvarado et al., 2015; Brian, 1974; R. Li et al., 2024; MacMillan et al., 2025; Rajakumar et al., 2012, 2018; Wheeler & Nijhout, 1981). While we have extensive knowledge of the role of JH in queen-worker (Brian, 1974; R. Li et al., 2024; Passera & Suzzoni, 1979) and worker-soldier (Rajakumar et al., 2018; Wheeler & Nijhout, 1981) development in ants, little is known of the role of ecdysone.

In the Florida Carpenter ant, *Camponotus floridanus,* we recently showed that JH could induce soldier development by a JH-mediated minor-soldier switchpoint in the last larval instar (MacMillan et al., 2025). This switchpoint is thermally-plastic and it was shown that developing soldiers differentially regulate the cellular reception and degradation of JH to enable their caste-specific development (MacMillan et al. 2025). At the phenotypic level, JH had a positive influence on head size, body size, head-to-body allometry, soldier caste identity, and it extended developmental timing (MacMillan et al., 2025). Here we use *C. floridanus* to investigate the role of ecdysone at the phenotypic and molecular level as well as its interaction with the JH signalling pathway during caste determination and differentiation. Specifically, we look at its role in caste-specific sizing and developmental timing as well as characterize caste-specific activity of ecdysone synthesizers, degraders, and the ecdysone signalling pathway. Furthermore, we investigated the interplay of JH and ecdysone signalling. We found that ecdysone has a negative role on sizing and blocks soldier development. Surprisingly ecdysone delays developmental timing and is negatively influenced by temperature. Unexpectedly, preliminary data suggests that in the context of JH and ecdysone both being high, there is an effect on sex such that females are masculinized, suggesting a role in sex differentiation.

## Results

### 20E regulates caste-specific sizing and developmental timing in *C. floridanus*

While JH has been shown to induce soldier development in *C. floridanus* (MacMillan et al., 2025), ecdysone and its role in caste development remains unexplored. To test the role of ecdysone during minor-soldier caste development, we treated bipotent larvae at the middle 4^th^ instar, which has been established as the sensitivity window when minor and soldier development diverge (MacMillan et al., 2025) and measured head size, body size, head-to-body allometry, and developmental timing. Compared to controls (EtOH), we found that head-to-body allometry was negatively influenced by 20E, the active form of ecdysone (Rewitz et al., 2006), where treated individuals exhibited a decrease in the slope of head size versus body size (Figure 1A; p=0.0474). Furthermore, head size (Figure 1B; p= 0.0357) and tibia length, which is an established proxy for body size (Teuscher et al., 2009) (Figure 1C; p= 0.0004) were significantly smaller than the control. Surprisingly, developmental timing was extended where the timing of cocoon formation (Figure 1D; p= 0.0009) and eclosure timing (Figure 1E; p= 0.0020) were both significantly delayed. In the ant *Myrm*ica *rubra*, ecdysone shortened developmental timing and supressed queen development (Brian, 1974) and soldier development in both *Pheidole* and *Camponotus* involves the extension of the last larval instar (MacMillan et al., 2025; Wheeler & Nijhout, 1981). Therefore, 20E is a negative regulator of soldier-like sizing and allometry, and it is unexpectedly capable of extending the last larval instar that is reflective of soldier development across ants.

**Figure 1:**
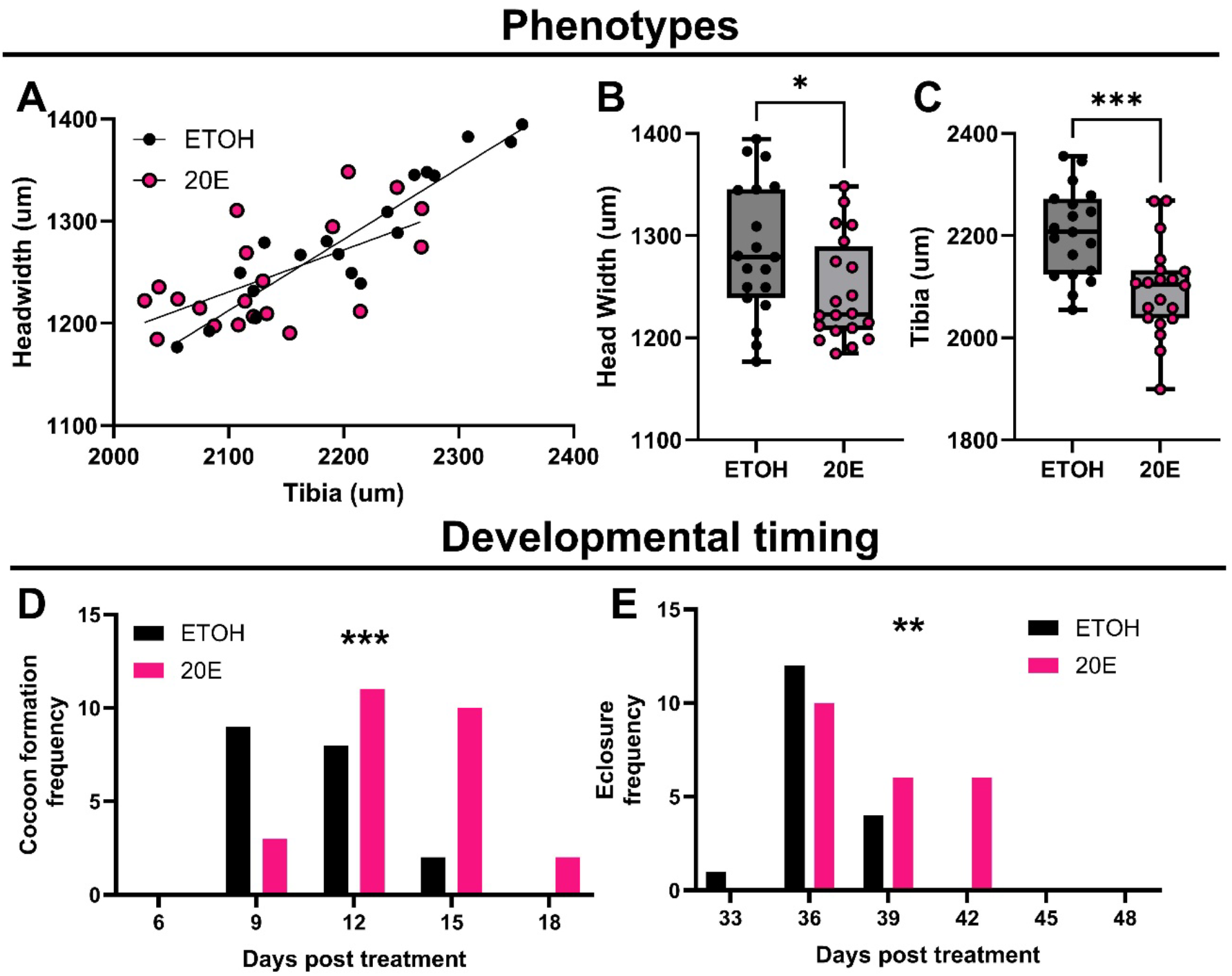
Role of ecdysone in sizing and developmental timing. Head-to-body allometry (A) was conducted using head width (B; p= 0.0357) and tibia length (C; p= 0.0004) to compare slopes of the control (ETOH) and treatment (20E) groups (A; p=0.0474). Developmental ting was determined using cocoon formation frequency (D; p= 0.0009) and eclosure frequency (E; p= 0.0020). The sample size for the control (black) was n=19 and the treatment (pink) was n=21.

### Molecular signatures of 20E signaling during caste determination

We next investigated the molecular basis of ecdysone signalling underlying the phenotypic changes we observed. In particular, we wanted to see whether ecdysone signalling is differentially regulated in a caste-specific manner at the level of synthesis, degradation, and/or reception. To do this, we looked at the relative expression of key ecdysone signalling pathway nodes at early (also known as ‘developing’) and late (also known as ‘terminal’) developing minor and soldier larvae. For synthesis we looked at the gene encoding Shade (*Shade*; also known as *CYP314a*), a cytochrome p450 gene that is the rate limiting (and last) step to synthesizing 20E, which is conserved across insects (Pan et al., 2021; Petryk et al., 2003; Warren et al., 2002). For degradation we looked at the gene encoding the cytochrome p450 enzyme CYP18a1 (*CYP18a1*), which is the anciently conserved degrader of 20E across arthropods (Schumann et al., 2018). For reception we looked at the genes encoding both nuclear receptors Ecdysone Receptor (*EcR*) and ultraspiracle (*usp*), the two of which form a heterodimer to mediate the tissue- and stage-specific expression of hundreds of genes downstream of the ecdysone signalling pathway and function similarly to type II nuclear receptors in mammals (Riddiford et al., 2000). Furthermore, *EcR* closely reflects 20E levels during development (Koelle et al., 1991) and *usp* is similar to the retinoid X receptor in mammals (Hall & Thummel, 1998; Oro et al., 1990; Yao et al., 1992). For downstream effectors of the pathway we looked at the gene that encodes Ecdysone induced protein 93 (*E93*) (Baehrecke & Thummel, 1995), Ecdysone induced protein 75 (*E75*) (Segraves & Hogness, 1990) and Broad *(br*) (Karim et al., 1993). All three of these factors are primary response factors found within the 20E signalling pathway, are conserved from crustaceans to insects, and are imperative for proper developmental transitions (Abdullah-Zawawi et al., 2021; Ishimaru et al., 2019; K. Li et al., 2015). For synthesis and degradation, we found that *Shade* and *CYP18a1*, respectively, expression was significantly increased in soldier larvae during early soldier differentiation as compared to early minor differentiation (Figure 2A; p<0.0001 and 2B; p=0.0073). While *Shade* remained increased during late soldier differentiation (Figure 2H; p=0.0012), *CYP18a1* was the same between late differentiation minors and soldiers (Figure 2I;). At the level of reception, we found a similar pattern where *ECR* and *usp* were both increased in expression during early soldier differentiation (Figure 2C; p= 0.0010 and 2D; p= 0.0004), but not late soldier differentiation (Figure 2J and 2K). At the effector level, *Br*, *E75*, and *E93* were all increased in expression during both early soldier differentiation (Figure 2E; p=0.0061, Figure 2F; p= 0.0001, and 2G; p<0.0001) and late soldier differentiation (Figure 2L; p=0.0274, Figure 2M; p=0.0034 and 2N; p=0.0062). Collectively these results suggest that all three regulatory levels (synthesis, degradation, and reception) as well as the downstream effectors, may be coordinated to enable soldier-specific differentiation. Furthermore, these results were unexpected as, it was previously shown that JH pathway components were elevated in soldier-destined larvae (MacMillan et al., 2025) and typically JH and 20E act antagonistically to regulate developmental transitions (Zhang et al., 2018). Therefore, these results are suggesting there may be simultaneous activity of both JH and ecdysone signalling during soldier differentiation.

**Figure 2:**
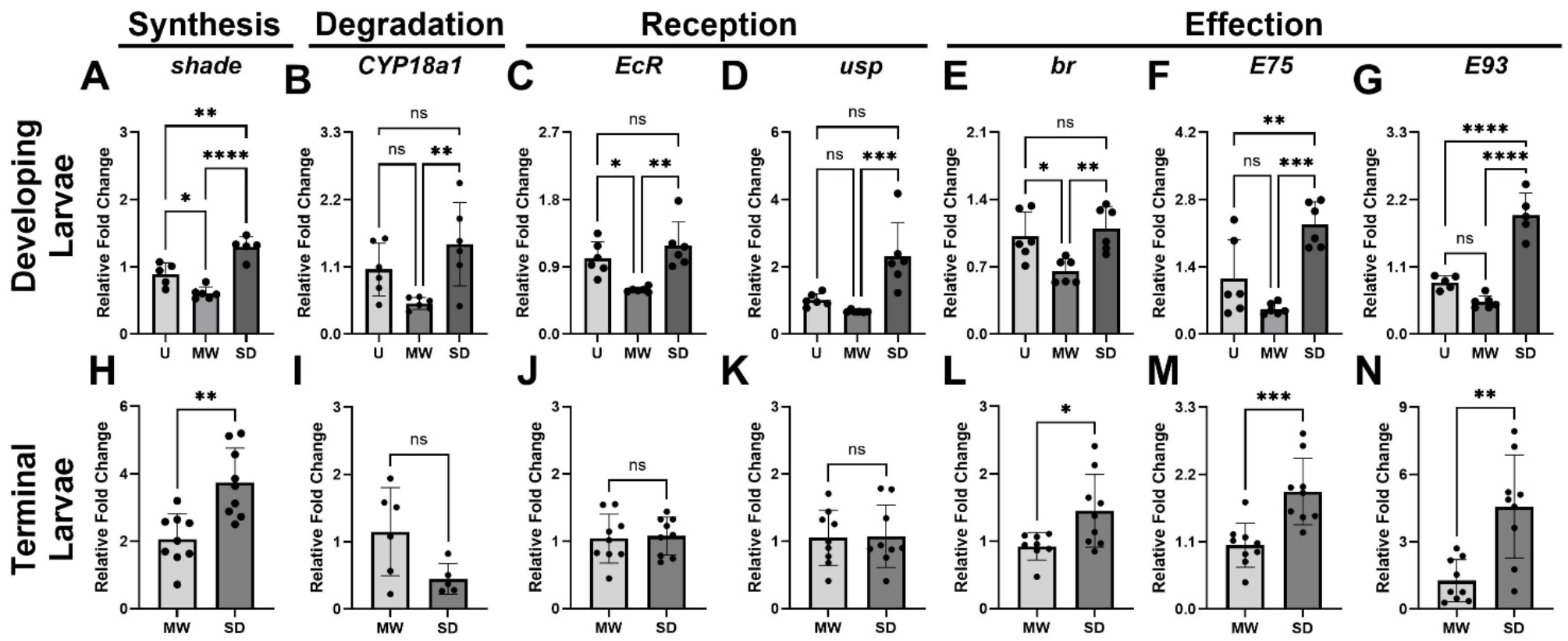
Molecular signatures of ecdysone during caste specific development. Relative expression levels of *Shade* (A, H), *CYP18a1* (B, I), *EcR* (C, J), *usp* (D, K), *Br* (E, L), *E75* (F, M), and *E93* (G, N) in *C. floridanus* developmentally (A-G) and terminally (H-N). Differences in expression between minors and soldiers developmentally (A<0.0001, B= 0.0073, C=0.0010, D= 0.0004, E= 0.0061, F= 0.0001, G<0.0001) and terminally (H=0.0012, I=0.0501, J=0.8202, K=0.9204, L=0.0274, M=0.0009, N=0.0023) in undifferentiated larvae (U), minors (MW) and soldiers (SD). The sample size was n=6-9 larvae.

### 20E & JH Crosstalk: Molecular Level

While surprising that ecdysone and JH signalling are both upregulated during soldier development, it has previously been shown that JH and 20E can both promote the same critical developmental targets (*Kr-h1*) in an array of insects (Depintor et al., 2025; Gospocic et al., 2021; Long et al., 2025). To characterize the interplay of the two pathways, and determine whether they synergize or antagonize one another, we examined the activity of a subset of our target genes (*EcR*, *usp*, *E93*) at two timepoints following hormonal manipulation of either JHA or 20E on bipotent larvae. Following the JHA treatment, we found that while *usp* did not change at the earlier timepoint (Figure 3A), it was downregulated following JHA compared to the control at the later timepoint (3B; p= 0.0022), and *EcR* did not differ between the control and JHA-treated at either timepoints (Figure 3C, 3D). Finally, *E93* was downregulated at the earlier timepoint compared to controls (Figure 3E; p= 0.0009), while, there was no difference at the later timepoint (Figure 3F). Therefore, this suggests that JH inhibits 20E signalling at both the reception and downstream effector levels.

**Figure 3:**
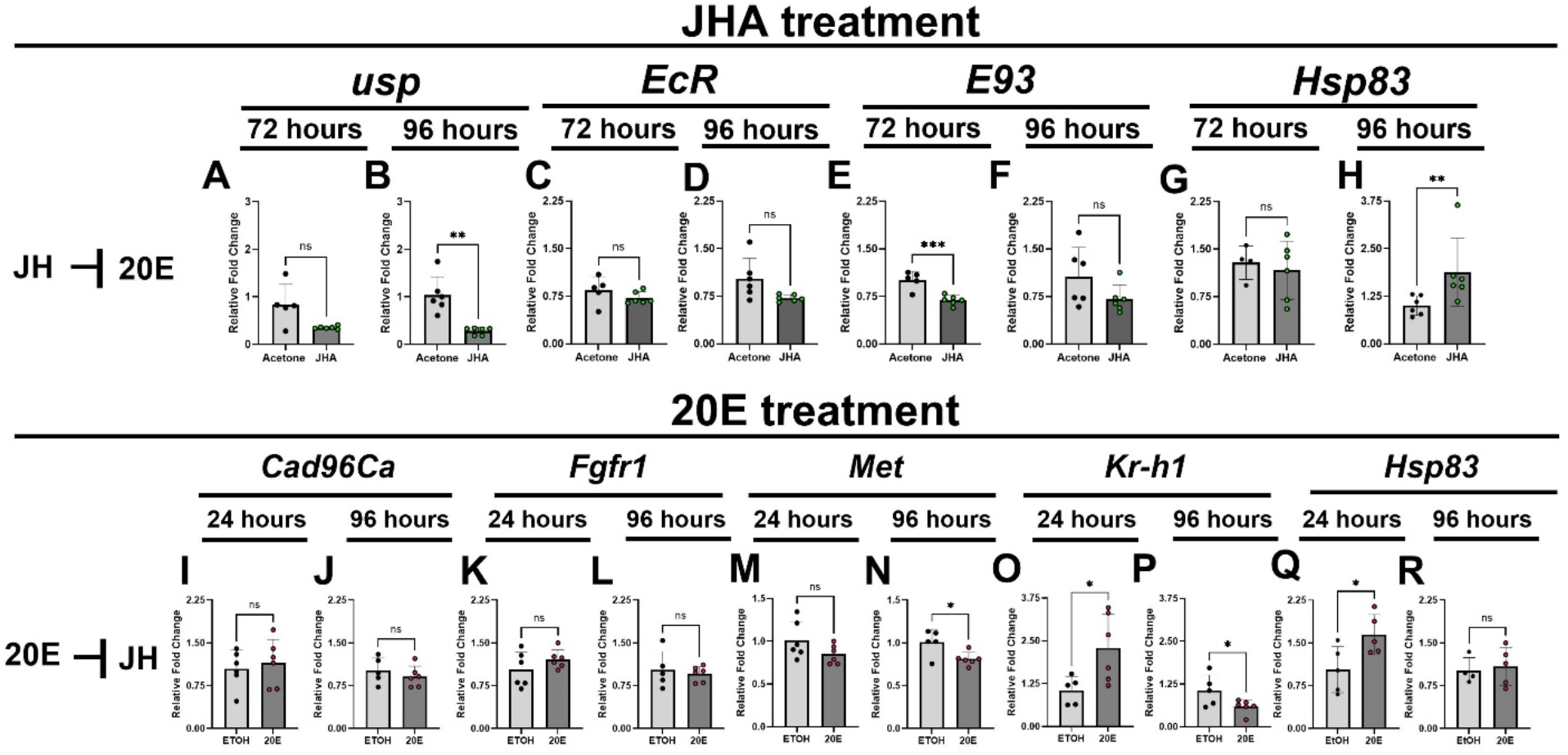
Molecular interplay between JH and ecdysone signaling. Relative expression levels of *usp* (A; p=0.0657, B; p= 0.0022), *EcR* (C; p=0.2504, D; p=0.0657), *E93* (E; p=0.0009, F; p= 0.1193), and *Hsp83* (G; p=0.6550 H; p=0.0087) following a 72-or 96-hour JHA treatment. Sample size for control (black) and JHA (green) was n=6. Relative expression levels of *Cas96Ca* (I; p=0.6273, J; p=0.4099), *Fgfr1* (K; p=0.2251, L; p=0.6281), *Met* (M; p=0.1285, N; p=0.0212), *Kr-h1* (O; p=0.0292, P; p=0.0483), and *Hsp83* (Q; p=0.0333, R; p=0.7076) following a 24-or 96-hours ecdysone treatment. Sample size for control (black) and 20E (pink) was n=6.

Reciprocally, we next wanted to characterize whether 20E has a positive or negative influence on the JH signalling pathway. For reception, we looked at the genes encoding Cadherin 96ca (*Cad96Ca*) and Fibroblast Growth Factor Receptor 1 (*Fgfr1*), which encode the two membrane receptors (Y.-X. Li et al., 2025), Methoprene-Tolerant (*Met*) which encodes the JH nuclear receptor (Jindra et al., 2015; Konopova & Jindra, 2007), and *Kr*-*h1* at two timepoints following 20E treatment. For *Cad96Ca* and *Fgfr1* we found that there was not a significant difference between the treatment and control at either timepoints (Figure 3I-L). For *Met* we found that while there was no significant difference at the earlier timepoint (Figure 3M), there was a decrease following 20E treatment at the later timepoint (Figure 3N; p= 0.0212). Finally, for *Kr-h1* we found that there was an increase at the earlier timepoint (Figure 3O; p= 0.0292) however, there was a significant decrease at the later timepoint (Figure 3P; p= 0.0483). Therefore, this data suggests that the reception and downstream, levels of the JH signaling pathway are mostly inhibited by 20E. In the case of *Kr-h1*, it is possible that the early increase is reflective of the positive transcriptional influence 20E can have through 20E response elements while the decrease that occurs later on is a reflection of 20E antagonizing JH-dependent regulation of *Kr-h1*. Overall, this suggests that there is generally an antagonism between these two pathways, yet the regulation of *Kr-h1* is a combination of cooperative and antagonistic crosstalk.

Finally, the heatshock protein 83 (HSP83; HSP90 in mammals) is known to be a critical mediator of ecdysone and JH signalling as a receptor complex chaperone (binding to MET in the JH receptor complex and USP in the ecdysone receptor complex), responsible for mediating the transcriptional activity of key downstream targets of both pathways, and transcriptionally-responsive to both ecdysone and JH (W. Liu et al., 2013). To understand the potential cooperative nature of the ecdysone and JH pathways, we treated bipotent larvae with either 20E or JHA and measured the expression of *hsp83*, the gene that encodes HSP83, at two timepoints. For JHA, *hsp83* expression did not change at the earlier timepoint (Figure 3G), but significantly increased at the later timepoint (Figure 3H; 0.0087) compared to controls. For 20E, *hsp83* expression is significantly increased at the earlier timepoint (Figure 3Q; p=0.0333), but not at the later timepoint (Figure 3R) compared to controls. It has previously been shown that *hsp83* is transcriptionally upregulated by both hormones in Lepidopteran *Helicoverpa armigera*, where it is upregulated earlier by 20E compared to JHA and then goes down in 20E, similar to what we have found (W. Liu et al., 2013). Altogether, *Kr-h1* and *hsp83* are two molecular levels that may facilitate the crosstalk between the ecdysone and JH pathways.

### 20E and JH Crosstalk: Phenotypic Level

Based on the natural expression of the ecdysone signalling pathway during minor and soldier differentiation and the crosstalk of 20E and JH at the molecular level (Figures 2 and 3), we next wanted to investigate the consequences of this crosstalk at the phenotypic level. While JH has been previously shown to generate larger individuals (MacMillan et al., 2025), here we show that 20E treatment leads to the development of smaller individuals. JH is typically topically applied and increases the head-to-body allometry of treated individuals (MacMillan et al., 2025), however 20E can only be microinjected and therefore we first microinjected JHA alone and assessed its effect on allometry. Compared to the control, we found that the slope of head-to-body allometry was significantly steeper in the JHA-treated samples (Figure 4A; p=0.0372), consistent with topical JHA. Based on the ability for microinjected JHA to increase head-to-body allometry, we then tested the effect of co-administering JHA and 20E on head-to-body allometry by co-injecting the two. Compared to controls, the co-treated individuals were the same as the control for their slope and y-intercept (Figure 4B). Therefore, while JH is capable of inducing an increase in head-to-body allometry akin to soldiers, ecdysone inhibits this capability, in a gatekeeping manner, when co-acting. Surprisingly, this treatment yielded an anomalous individual (1/22; Figure 4C) that developed a third eye (Figure 4D) and this supernumerary eye exhibited male-like eye morphology. This anomaly suggests that beyond their opposing roles in sizing and worker-soldier caste development, the interaction between JH and ecdysone may have sex-inducing properties. To further test this hypothesis, we collected soldier-destined individuals that have naturally high JH signaling activity and injected them with either 20E or control. While controls all developed into soldiers, which are females, one soldier-destined 20E-injected anomalous individual (1/11; Figure 4E) surprisingly developed into an adult that appears male-like (Figure 4F) when comparing to a natural male (Figure 4G). Unexpectedly, this male-like individual possesses external genitalia (Figure 4H left) similar to that of a typical female solider (Figure 4H right), that lacks the external genitalia of males including the aedeagus (Figure 4H bottom). This is unexpected because the lab colonies used never produce males, whether in the colony or in experimental replicates. Altogether, this very preliminary data suggests that through some unknown mechanism, JH and ecdysone are together capable of influencing sex determination in *C. floridanus*.

**Figure 4:**
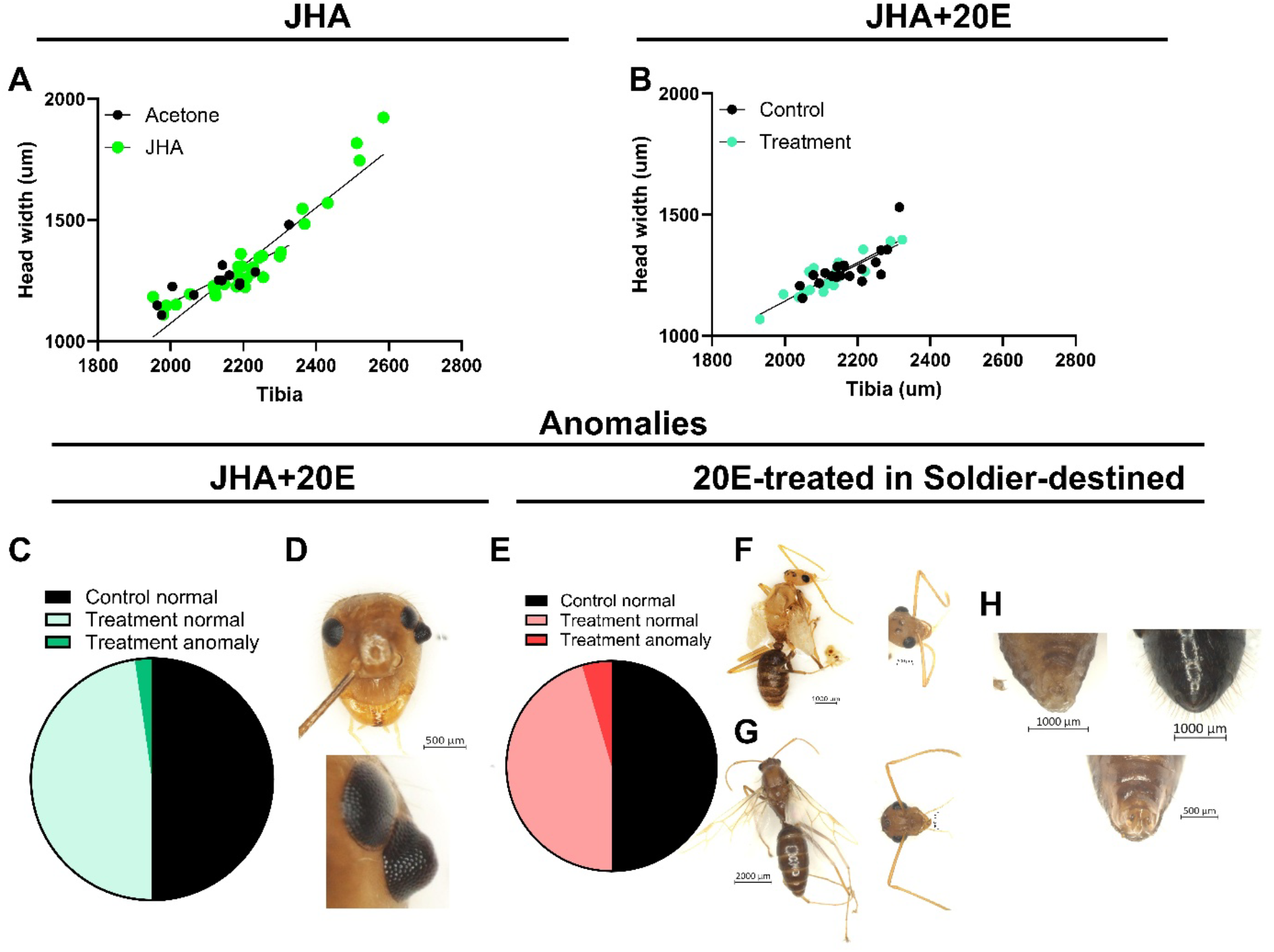
Antagonism and synergy of JH and ecdysone in sizing and developmental timing. Head-to-body allometry to compare the slope (A; p=0.0372) of JH injected individuals, control n=10, treatment n=31. Head-to-body allometry to compare the slope (B; p= 0.7451) and y-intercept (B; p= 0.7134) between JHA + 20E treatment and acetone + ETOH control, n=22 for both control and treatment. Percentage of anomalies produced in JHA + 20E treatments was 4.55% (C) and produced a female individual with a 3^rd^ eye; the eye had male morphology, a closeup below (D) (left antenna removed for morphometrics), the sample size was n=22 for both treatment and control. Percent of anomalies produced in 20E soldier-destined treatments was 9.09% (E). An individual with male morphology (F; whole body and head), compared to a natural male (G; whole body and head); the sample size was n=11 for both treatment and control. Abdomen from ventral view showing the external genitalia region (H) of individual from (F) to the left, a natural soldier to the right, and natural male below.

### Thermal Plasticity of 20E

Previously it was shown that the minor-soldier JH-dependent switchpoint and JH signalling are further sensitized by increasing temperature, where temperature induced both large soldiers and a higher frequency of soldiers (MacMillan et al., 2025). Therefore, we wanted to investigate whether temperature decreases ecdysone signalling in parallel or influences size through JH independent of ecdysone signalling. To test this, we reared bipotential larvae at 22° C or 27° C and we measured relative expression of the downstream effector *E93*. We found that individuals developing at 27° C exhibited a significant reduction in expression of *E93* relative to those developing at 22° C (Supplemental figure 1A; p=0.0335). Therefore, temperature has opposite effects on JH compared to ecdysone and the inhibiting influence of ecdysone on both head-to-body allometry and the minor-soldier switchpoint, may be negated at higher temperatures. To further test this, we conducted a co-injection of JHA and 20E and raised the individuals at 27°C. We found that the slope of head-to-body allometry was increased in the treatment relative to the control (Supplemental figure 1B; p=0.0067), resembling more soldier-like allometry and therefore blocking the gatekeeping effect of ecdysone on head-to-body allometry. Finally, in the context of temperature having diverging effects on JH and ecdysone and the crosstalk between JH and ecdysone converging at both *Kr-h1* and *Hsp83*, we wanted to further examine the influence of temperature on this crosstalk. It was previously shown that *Kr-h1* responds positively to increased temperature, therefore we investigated how *Hsp83* expression changes with temperature. We found that bipotent individuals developing at 27° C exhibited a significant reduction in expression of *Hsp83* relative to those developing at 22° C (Supplemental figure 1C; p=0.0173). Collectively, temperature positively regulates JH signalling and the JH-ecdysone crosstalk gene *Kr-h1*, but negatively regulates ecdysone signalling and the JH-ecdysone crosstalk gene *Hsp83*. Therefore, increasing temperature may coordinate JH-ecdysone crosstalk to enable soldier development by both: increasing *Kr-h1*, enhancing *Kr-h1* in mediating downstream processes of JH-dependent soldier development, and decreasing *hsp83*, blocking the receptor function of *hsp83* involved in 20E’s role as a negative gatekeeper of soldier development.

## Discussion

### 20-hydroxyecdysone as both a negative regulator of size and brake on soldier development

Juvenile hormone and its signalling pathway has been shown to positively regulate soldier-specific caste determination and differentiation (Fig. 5A-C) (MacMillan et al., 2025). Our data show that ecdysone is a negative regulator of growth in *C. floridanus* and that the JH and ecdysone signalling pathways antagonize each other. This is consistent with that typically found across insects (S. Liu et al., 2018). Furthermore, while JH signaling has been previously established to be positively regulated by temperature in *C. floridanus* (MacMillan et al., 2025), here we show that ecdysone signalling is negatively regulated by temperature. Additionally, while these hormones block each other when co-administered at a baseline temperature, at a high temperature the influence of JH overcomes that of ecdysone, leading to a size increase. The inverse relationship of temperature and ecdysone during *C. floridanus* caste development is in contrast to the positive relationship found during the plastic development of eyespot size and number in butterflies and size plasticity in fire ants (Bahramifarid et al., 2026; Brakefield et al., 1996, 1998; Koch et al., 1996). Unexpectedly, we found that ecdysone synthesis and reception is higher in developing soldiers and that ecdysone can increase *Kr-h1* expression similarly to what has been shown with JH (MacMillan et al., 2025). JH and 20E are also known to positively regulate *Kr-h1* in the context of fly and honeybee development, and behavioural plasticity in ants (Depintor et al., 2025; Gospocic et al., 2021; Long et al., 2025). In particular, it has been shown that the cis-regulatory region of *Kr-h1* has both ecdysone and JH response enhancer elements (Long et al., 2025). In termites, alongside JH reception, the ecdysone receptor EcR is required for proper soldier differentiation (Masuoka & Maekawa, 2016). Several potential mechanisms could reconcile their being both antagonistic and synergistic interactions between these two pathways. *Kr-h1* is a transcription factor that can activate or suppress transcription of target genes (Ojani et al., 2018) and that 20E and JH may be providing different regulatory inputs that change Kr-h1 function by influencing chromatin accessibility (Long et al., 2025) or post-transcriptional modifications of *Kr-h1* (Pecasse et al., 2000) or co-factors interacting with Kr-h1. Altogether, we propose that in *C. floridanus*, while JH induces soldier development, ecdysone blocks soldier development in a gatekeeper fashion and may further be differentially regulated during soldier differentiation to control intracaste soldier sizing (Fig. 5A). More generally, 20E could potentially play a role in organ-specific and systemic regulation of developmentally plastic growth across insects.

**Figure 5:**
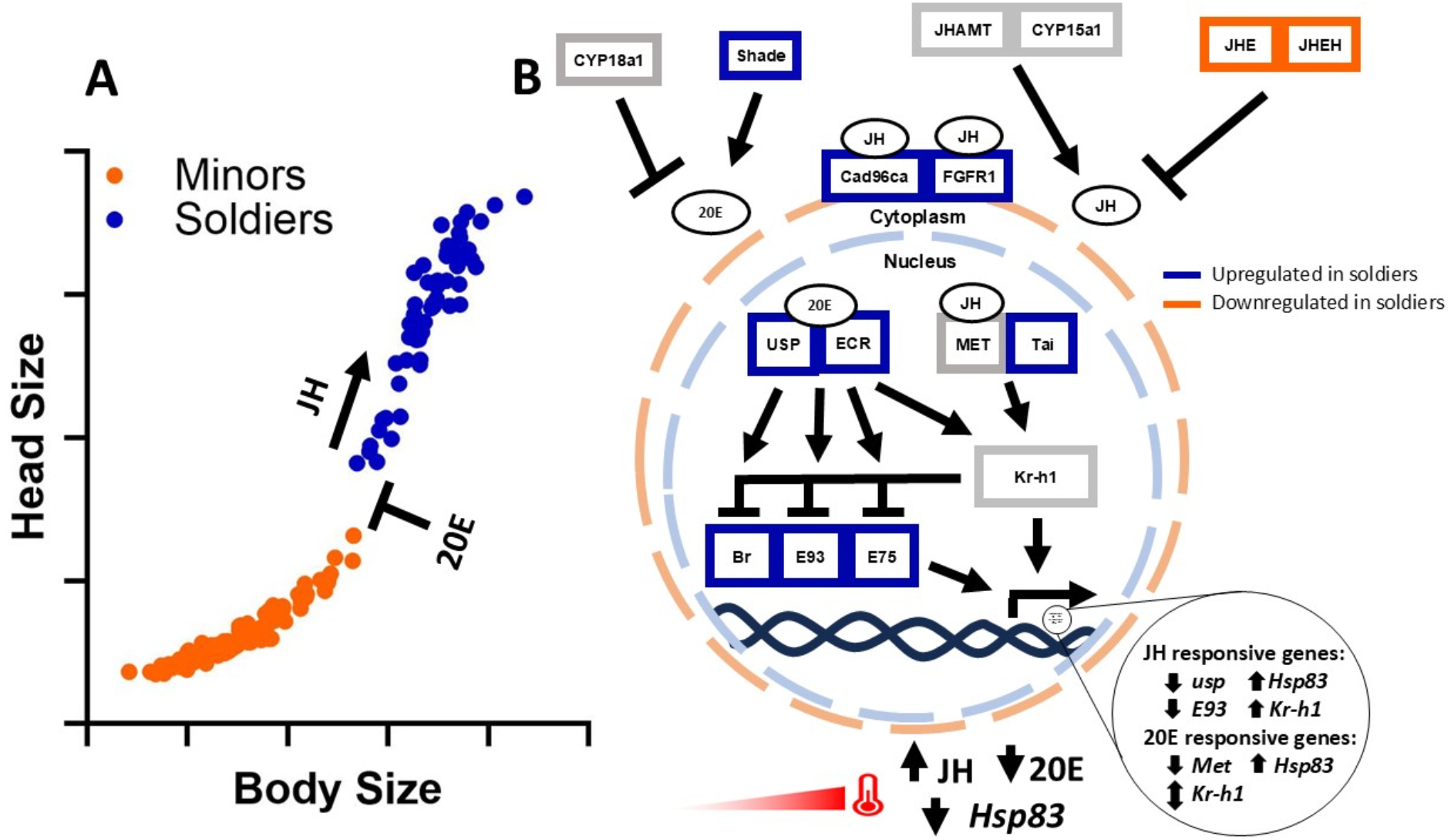
The complex interactions of JH and 20E in relation to size, temperature, and caste. In the context of natural head-to-body size variation across a *C. floridanus* colony (modified from Alvarado et al.., (2015)), our results suggest that JH is imperative for soldier development and that 20E may act in an antagonistic way to prevent soldier development (A). JH and 20E signaling pathway components are represented (B) synthesizers; *JHAMT*, *CYP15a1*, and *Shade*, degraders; *JHE*, *JHEH*, and *CYP18a1*, receptors; *Cad96Ca*, *Fgfr1*, *Met*, *Tai*, *usp* and, *EcR*, and downstream effectors; *Kr-h1*, *br*, *E93*, and *E75*. Based on our findings alongside those of MacMillan et. Al., (2025), genes highlighted in blue are upregulated during soldier development and genes highlighted in orange are downregulated during soldier development. Temperature on JH has a positive relationship whereas, temperature on 20E and *Hsp83* have a negative relationship. Genes responding to JHA or 20E treatments are indicated (B).

### 20E & JH: blurring the line between sexual dimorphism and morph-specific plasticity

In horned beetles, alongside the differences in males and females, there is phenotypic plasticity within the males such that males experiencing a juvenile hormone pulse leads to the development of large horns and large bodies, while males that do not experience this JH pulse do not grow these large horns and appear more female-like in regards to the horns (Emlen & Nijhout, 1999). Before the stage that this JH switch occurs, there is an earlier period of development where there is a pulse of ecdysone in females and hornless males, but not in males who will eventually grow large horns. This earlier pulse of ecdysone may play a key role in inducing female development, including feminizing eventual hornless males. In our case, JH similarly influence size and head-to-body allometry, inducing soldiers that are reminiscent of the large horned males in beetles (Emlen & Nijhout, 1999). In contrast, rather than having a feminizing effect, we observed that ecdysone may have a masculinizing effect in *C. floridanus* such that it can induce male morphologies individuals that were treated with JH and ecdysone at the minor-soldier switchpoint or injecting soldier-destined individuals that have high endogenous levels of JH, with 20E. This inverse relationship of 20E and feminizing versus masculinizing individuals in beetles and ants is further reflected in the fact that beetles exhibit male plasticity while ants exhibit female plasticity. Future work aimed at investigating how 20E, along with its relationship with JH, can influence sex and sexual differentiation.

### Connecting sex and growth differentiation: Morphogens, sex and non-sex steroid hormones

In vertebrates, sexual differentiation is regulated by sex steroid hormones including androgens and estrogens (Arnold, 2009; Elmelund et al., 2026; Nagahama et al., 2021). Beyond sex steroid hormones, there are molecules that typically regulate morphogenesis and development that are also known to influence sex. Specifically, retinoic acid produced by the mesonephros of the embryo during sexual differentiation of the primordial germ cells (PGC), is differentially regulated by the retinoic acid degrader cytochrome p450 26b1 (Cyp26b1), which controls the timing of meiosis and sexual identity of the PGCs (Bowles et al., 2006; Koubova et al., 2006). This example of a hormone degrader playing a vital role in sex identity is reminiscent of our findings that the cytochrome p450 family Cyp18a1, the 20E degrader, being increased in expression during *C. floridanus* soldier development. Furthermore, sex steroid hormones are known to influence non-sex trait variation including appendage size. For example, in mammals, the ratio between the lengths of the 2^nd^ digit (2D) and 4^th^ digit (4D) differs between males and females where males have a 2D:4D ratio <1 and females are ≥ 1 and this allometry is controlled by the balance of the sex steroid hormones androgen and estrogen (Zheng & Cohn, 2011). Finally other steroid hormones in mammals can influence trait and body size, including vitamin D and its degrader Cyp24a1 regulating heart size and glucocorticoid signalling regulating body size (Engblom et al., 2007; Han et al., 2019; Tronche et al., 2004). Our findings suggest that the interaction of JH and 20E may have a sex identity role alongside their plasticity role in caste identity, and we propose that future work aims to disentangle the blur between sex and size plasticity and their developmentally dimorphic regulation.

## Methods

### Ant husbandry

Ant colonies were collected from Gainesville, Florida and were kept in an incubator at 25° C with 60% humidity and a 12-hour light dark cycle. Colonies had a constant supply of water and sugar water. Colonies were fed 3-4 times a week with Bhatkar & Whitcomb diet (Bhatkar & Whitcomb, 1970) and were fed mealworms twice a week.

### Developmental stages characterized and manipulated

Larvae used for experiments were selected from the same criteria indicated in MacMillan et al. 2025, where bipotential individuals were 3.5-4.2mm, soldier-destined larvae were 6.1+mm, undifferentiated larvae were 2.8-3.2mm, and developing minor larvae were 3.8-4.2mm. All of these larvae types had a brown gut to indicate ongoing development. Terminal minor larvae were 4.7-5.5mm and terminal soldier larvae were 6.7+mm both had a black gut to indicate the end of larval development (MacMillan et al., 2025).

### 20E treatments

Individual bipotent larvae or soldier-destined larvae were microinjected with either 100ng/ul of 20E or 9.9% ethanol (control) using an Eppendorf CellTram 4r (oil) microinjector, controlled by a Narishige micromanipulator with the aid of a Zeiss SteREO Discovery.V8 microscope. These individuals were raised in replicates composed of a 2 minor worker adults : 1larva ratio and were fed 3 times a week with mealworms and the Bhatkar & Whitcomb diet (Bhatkar & Whitcomb, 1970). Microinjection needles were made using a Sutter Instrument P-97 needle puller. Environmental conditions for the experiments are the same as colonies described above. Individual larvae collected for RT-qPCR analysis were collected after 24- and 96-hour time points and placed in the -80°C freezer. Individuals collected for phenotypic analysis were collected upon eclosure and preserved in 70% ethanol for dissection and microscopy.

### JH treatment

Individual bipotential larvae were microinjected (as above) or topically treated (MacMillan et al., 2025) with 5mg/ml methoprene (a synthetic JH analog; Sigma) or acetone (control). These individuals were raised similarly to the ecdysone treatments and the needles were pulled in the same manner. Individual larvae collected for RT-qPCR analysis were collected after 72- and 96-hour time points and placed into individual tubes and kept in the -80°C freezer. Individuals were collected for phenotypic analysis were collected upon eclosure and preserved in 70% ethanol for dissection and microscopy.

### JH + 20E co-injections

Bipotential larvae were microinjected with 100ng/ul of 20E + 5mg/ml methoprene (treatment) or 9.90% ethanol + 9.95% acetone (control). Individuals were treated and raised as above. Environmental conditions for the experiments are the same as indicated above and were reared in either 25°C or 27° C. Individual larvae were collected for RT-qPCR analysis were collected after 24- and 96-hour time points and placed into individual tubes and kept in the -80°C freezer. Individuals collected for phenotypic analysis were collected upon eclosure and preserved in 70% ethanol for dissection and microscopy.

### Temperature treatment

Individual bipotent larvae were collected and placed at either 22°C or 27°C and were raised and fed as above. Individual larvae were collected for RT-qPCR analysis were collected after 96-hours after the treatment and were placed into individual tubes and kept in the -80°C freezer to be used for RNA extraction.

### Microscopy and morphometrics

Adult sample head width and tibia lengths were measured using the ZEN Pro 3.2 Software at 25x magnification using a Zeiss Axio Zoom.V16 stereomicroscope. Head width is the hallmark measurement used to differentiate castes in *Camponotus floridanus* (Alvarado et al., 2015; Diniz-Filho et al., 1994a; MacMillan et al., 2025) and tibia length is an extensively used proxy for body size in *Camponotus* (Teuscher et al., 2009) and other ants (Diniz-Filho et al., 1994b; Kaspari & Weiser, 1999).

### Developmental timing

Developmental timing was recorded by daily observations of developing individuals to track the timing of cocoon formation and eclosure.

### RNA extraction & cDNA synthesis

RNA and cDNA synthesis was conducted following what was described in MacMillan et al, 2025. Briefly, samples were removed from the -80° freezer and total RNA exaction and purification was performed using a method combining the traditional Trizol extraction with the Direct-zol RNA MicroPrep kit (Zymo Research). Turbo DNase (Invitrogen) treatment was used to remove residual genomic DNA from the RNA samples and then cDNA was synthesized using the SuperScriptIV First-strand Synthesis System kit from Invitrogen. RNA and cDNA used a 1/20 dilution as the working concentration for RT-qPCR (MacMillan et al., 2025).

### RT-qPCR

RT-qPCR was performed to determine relative expression of key genes in the JH and 20E pathway and was conducted following what was described in MacMillan et al, 2025. JH-related genes were investigated using previously described primers (MacMillan et al., 2025). RT-qPCR primer pairs including at least one primer that spanned across an exon-exon boundary (Table S1) and RT-qPCR was conducted with SSO advanced SYBR green (BioRad) and ran on a C1000 CFX96 machine (BioRad). The ΔΔCT (Livak & Schmittgen, 2001) method was used to calculate relative fold changes and the gene *RpL32* (also known as rp49) was used as the reference gene, which is an extensively used housekeeping gene amongst insects as it remains stable across hormonal manipulations and developmental changes (Alvarado et al., 2015; Colombani et al., 2012; Depintor et al., 2025; Gospocic et al., 2021; MacMillan et al., 2025).

### Statistical analysis

Graphpad Prism was the primary application used for statistical analysis. RT-qPCR data took into account normal distribution and equal variance for the analysis. When a 2-group comparison was done; a Shapiro-Wilk test for normality was conducted and a F-test was done to rule out unequal variance. If data was normally distributed with equal variance, an unpaired t-test was conducted. If the data was normally distributed with unequal variance a Welch’s test was performed. If the data was not normally distributed, a Mann-Whitney test was done. When a 3-group comparison was done; the Shapiro-Wilk test and Browns-Forsythe test were conducted to assess normality and variance. If the data was normally distributed with equal variance, an ordinary one-way ANOVA was done. If the data was normally distributed with unequal variance a Welch’s one-way ANOVA was conducted. If the data was not normally distributed, a nonparametric ANOVA was performed. All statistical tests for RT-qPCR analysis used a p-value <0.05.

For phenotypic analysis, a simple linear regression with an ANCOVA was used to determine statistical significance between treatment and control groups. The slope represents the head-to-body allometric relationship where an increase signifies disproportionate head-to-body growth (and vice versa for decrease). The y-intercept also reflects this allometric relationship where an increase signifies an increase in head size relative to body size across groups (and vice versa for decrease). Statistical analysis of head widths, tibia lengths, and developmental timing between treatment and control groups were determined using the same method as the 2-group comparison explained above.

## Acknowledgements

We thank L. Davis and the Abouheif lab for help with ant collection. We also thank A. Rajakumar for comments and Rajakumar Lab members for both comments and husbandry support. We acknowledge the support of the Natural Sciences and Engineering Research Council of Canada (NSERC) [RGPAS-2021-00006; RGPIN-2021-04399; RTI-2021-00710]; Canada Foundation for Innovation and the Ontario Research Fund [40142].

## Author Contributions

Conceptualization, O.M. and R.R.; Data curation, O.M.; Formal analysis, O.M, R.R.; Funding acquisition, R.R.; Investigation, O.M., A.C., C.G., J.S., and D.Q.; Methodology, O.M. and R.R; Projection administration, R.R.; Resources, R.R.; Supervision, R.R.; Validation, O.M.; Visualization, O.M. and R.R.; writing—original draft, O.M. and R.R.; writing—review & editing, O.M. and R.R.

## Supplemental Figure and Table

**Supplemental Figure 1:**
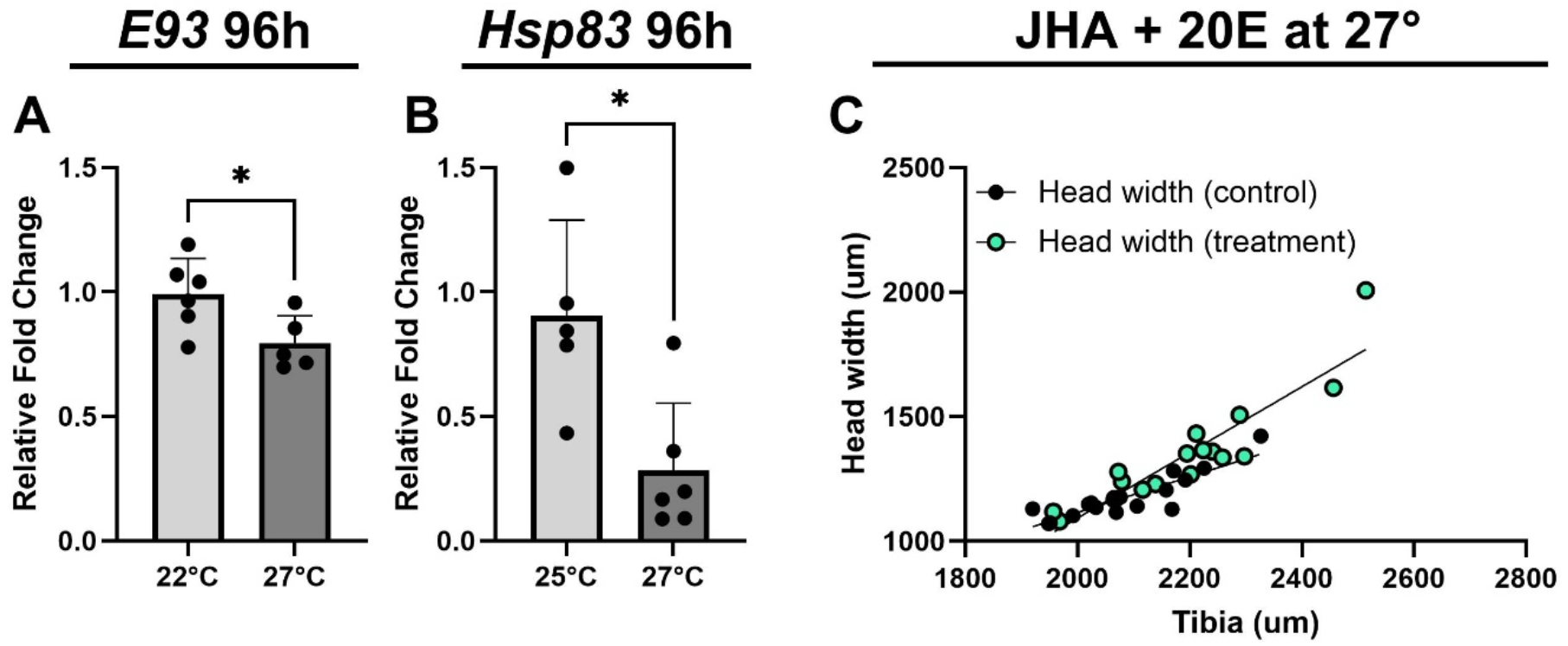
Ecdysone is negatively regulated by temperature preventing antagonism of positive temperature-dependent JH regulation of size. Relative expression of *E93* (A; p=0.0335), and *Hsp83* (B; p=0.0173) following a 96-hour treatment of either 22° or 27°C, the sample size was n=6. Head-to-body allometry was conducted to compare the slope (B; p=0.0067) of JHA+20E treatment compared to control samples, n=18 for control, and n=18 for treatment.

**Table S1:**
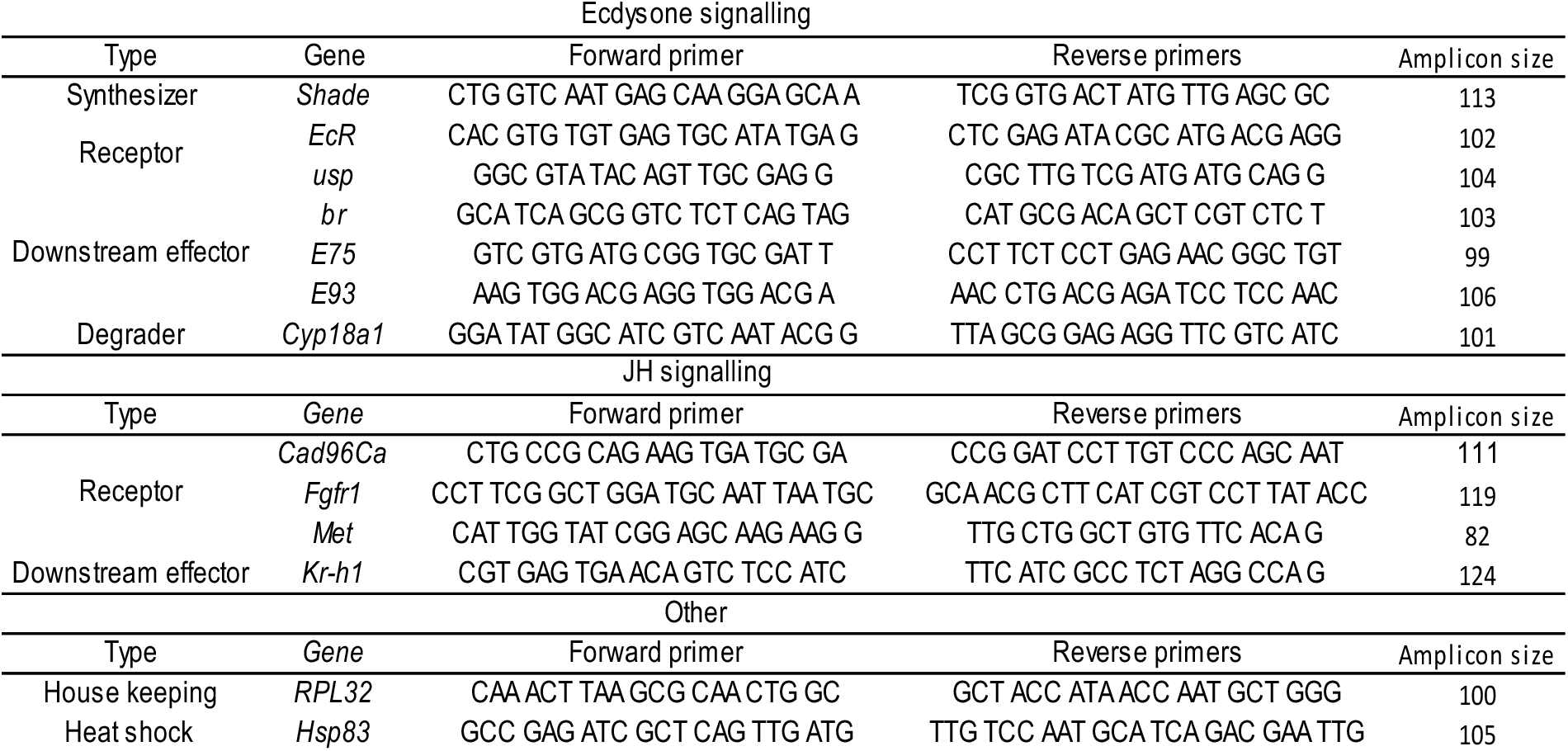
qPCR primes used for JH and ecdysone signalling pathways to determine relative gene expression. . Primer sequences and amplicon length are present, and one primer from each gene spanned an exon-exon junction.

## Notes

### Competing Interest Statement

The authors have declared no competing interest.

